# Evolution into chaos – implications of the trade-off between transmissibility and immune evasion

**DOI:** 10.1101/2024.06.29.601333

**Authors:** Golsa Sayyar, Ábel Garab, Gergely Röst

## Abstract

Predicting viral evolution presents a significant challenge and is a critical public health priority. In response to this challenge, we develop a novel model for viral evolution that considers a trade-off between immunity evasion and transmissibility. The model selects for a new strain with the highest invasion fitness, taking into account this trade-off. When the dominant strain of the pathogen is highly transmissible, evolution tends to favor immune evasion, whereas for less contagious strains the direction of evolution leads toward increasing transmissibility. Assuming a linear functional form of this trade-off, we can express the long-term evolutionary patterns following the emergence of subsequent strains by a non-linear difference equation. We provide sufficient criteria for when evolution converges, and successive strains exhibit similar transmissibility. We also identify scenarios characterized by a two-periodic pattern in upcoming strains, indicating a situation where a highly transmissible but not immune-evasive strain is replaced by a less transmissible but highly immune-evasive strain, and vice versa, creating a cyclic pattern. Finally, we show that under certain conditions, viral evolution becomes chaotic and thus future transmissibilites become unpredictable in the long run. Visualization via bifurcation diagrams elucidates our analytical findings, revealing complex dynamic behaviors that include the presence of multiple periodic solutions and extend to chaotic regimes. Our analysis provides valuable insights into the complexities of viral evolution in the light of the trade-off between immune evasion and transmissibility.

## 1. Introduction

Predicting the evolution of viruses is both a significant challenge and a major public health concern. Numerous studies have investigated virus evolution by introducing a trade-off between pathogens’ epidemiological traits (virulence, transmissibility, and clearance). However, the majority of these investigations have primarily focused on the trade-off between virulence and the transmissibility. Anderson and May [5] argued that a parasite cannot increase its transmission rate without shortening its infectious period by harming its host, and they showed that the parasite should adopt an optimum intermediate level of virulence. This trade-off approach inspired many subsequent works. This phenomenon is explored in detail in Chapter 11 of the book “Evolutionary Dynamics” [23], where the epidemiological dynamics of at least two parasite strains competing for the same host have been studied. If transmissibility is proportional to virulence, the basic reproduction number is an increasing function of virulence, and selection favors strains with higher virulence and thus higher infectivity. However, when transmissibility is a saturating non-linear function of virulence, the basic reproduction number becomes a hump-shaped function of virulence, maximized at an optimal intermediate level. Therefore, if the virulence of a parasite population is greater than this value, then selection will reduce virulence; otherwise, it will increase it.

The study [3] investigated the emergence of a convex trade-off between transmission and virulence, utilizing a model that explicitly incorporates within-host dynamics. Their analysis indicated the robustness of this convex trade-off. Additionally, they demonstrated that small variations among parasites or hosts might blur the trade-off curve, as parasites with the same within-host growth rate can express different virulence or transmission values for each host they infect. Further in-depth discussion of the transmission– virulence trade-off can be found in [1, 2, 19, 26] and references thereof.

Other models have been developed that incorporate trade-offs between additional traits. For instance, [18] modeled an inherent trade-off between contact and transmissibility using a basic susceptible–infected–recovered (SIR) framework by differentiating between mild and severe infections to indicate that increasing symptom severity will tend to decrease contact rates and increase the probability of transmission given contact. The work [6] studied the host-pathogen interaction for the case in which hosts may become at most doubly infected. It was discovered that in the presence of frequent double infections, increased virulence is favored; but when pathogens become more virulent, the force of infection will decrease, followed by lower virulence again. They concluded that the endemic steady state of the virulence depends on the interaction within hosts as well as on the interaction at the population level.

On 12 March 2020, the World Health Organization (WHO) declared a new worldwide pandemic, COVID-19, which originated in China in December 2019 and rapidly spread around the world, causing millions of deaths [22]. The most common SARS-CoV-2 variants of concern, including the wild type, Alpha, Beta, Gamma, Delta, and Omicron, show great diversity in their transmissibility and virulence [16, 21, 29].

This diversity, and the need to assess the evolutionary trajectory of new variants, urged scientists to re-examine the potential trade-off between virulence and transmissibility of SARS-CoV-2 [15]. Other studies of SARS-CoV-2 evolution explore how different vaccination strategies could impact infection dynamics and antigenic evolution in partially immune populations [27], or discuss the impacts of immune escape and transmissibility on the endemic load of SARS-CoV-2 [24]. A major concern related to new variants was their ability to evade immunity and its potential impact on the severity of upcoming waves [7, 28, 31]. According to [10], antibody resistance may compromise viral fitness, such as in the B.1.351 variant, which resists neutralization by antibodies but also loses enhanced transmissibility as a consequence of the immune-escape mutations. The JN.1 variant showed higher immune evasion compared with BA.2.86 and other strains, at the expense of reduced human ACE2 binding, and this evolutionary pattern has been observed in the previous transition from BA.2.75 to XBB [32]. Hence, it is natural to consider a potential trade-off between immune evasion and transmissibility. Such a trade-off is further supported by the analysis published in [25].

While the classical trade-offs have been extensively investigated, there remains a significant gap in mathematical understanding of the implications of a trade-off between transmissibility and immune evasion. Viral evolution often operates under selective pressures that favor maximizing the basic reproduction number *R*_0_, which measures the average number of secondary infections caused by a single infected individual in a fully susceptible population. In our framework, however, optimal evolutionary strategy depends on the resident strain and refers to the evolutionary pathway that maximizes the invasion fitness of a viral strain, balancing the trade-off between immune evasion and transmissibility. Understanding this balance is crucial for predicting the long-term behavior of pathogens and developing effective public health interventions.

To account for such a trade-off in viral evolution, we construct a novel evolutionary model by the following steps (Fig. 1.1):

**Figure 1.1:**
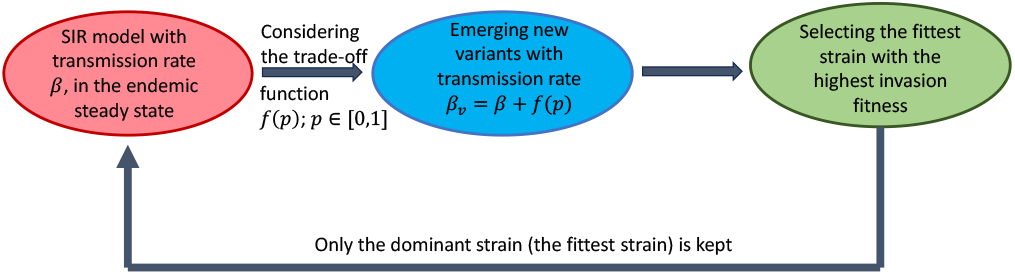
The model outline flowchart: Beginning with the SIR model with a transmission rate *β*, while the system is in an endemic steady state (red). New strains with immune-evasive property *p*, where *p* ∈ [0, 1], emerge. Assuming a trade-off between transmissibility and immune evasion, denoted by *f* (*p*), the transmission rates of these new strains are *β*_*v*_ = *β* + *f* (*p*) (blue). As the evolution tends to favor the strain with maximal invasion fitness (maximizing the invasion reproduction number), we consider the newly emerging strain which is the fittest with a transmission rate *β*_*v*_ = *β* + *f* (*p*^max^) (green). The system transitions to a new endemic steady state characterized by a novel transmission rate determined by the trade-off function. The process returns to the initial step and the evolutionary process is continued.

1. We initiate our analysis by investigating an SIR system with a single strain in an endemic steady state.
2. Upon the emergence of new strains distinct from the previous strain, we introduce a parameter *p* ∈ [0, 1] to denote their immune-evasive property, which represents the fraction of recovered individuals from the resident strain that can be infected by the new strain. Strains with *p* = 0 are unable to infect anyone with prior immunity, while strains with *p* = 1 can infect all individuals. Additionally, we assume a trade-off between transmissibility and immune evasion, making the transmission rate of new strains dependent on *p*. Subsequently, our focus shifts to the introduction of novel strains, with the aim of determining whether their emergence is attributed to their ability to evade immunity or their heightened transmissibility.
3. Next, we select the fittest strain by maximizing the invasion fitness, and the system goes to the new endemic steady state characterized by this new strain with a new transmission rate determined by the trade-off function.
4. Return to Step 1 and continue the evolutionary process.

This evolutionary process is depicted in Fig. 1.1. Although many adaptive dynamics frameworks emphasize the role of small mutational steps in evolutionary processes [8], our model distinguishes itself by allowing larger evolutionary jumps with significant differences between resident and invader strains. This reflects real-world observations such as the COVID-19 pandemic, where, from the perspective of a given host population, the evolution was characterized by the emergence of vastly different strains with the ability of replacing the resident strain without a gradual sequence of mutations (which may have occurred elsewhere). An important aspect of our model is the difference between the epidemiological dynamics of the resident and invader strains. Specifically, the resident strain cannot infect recovered individuals due to acquired immunity. In contrast, invader strains may exhibit varying degrees of immune evasion, which allows them to re-infect a part of the recovered population. This asymmetry is important for shaping the invasion fitness of new strains and influences the direction of their evolutionary trajectory.

Through our theoretical framework, we shall investigate whether in the short term such evolution points toward enhanced transmissibility or enhanced immune evasion (Theorem 2.1). We analyze the transmissibility and the immune evasion capability of successive invading strains, and explore this evolution over the long term as well. We provide explicit conditions when the evolution converges to a certain transmissibility (Theorem 3.2), but we also find situations where there exists an alternating pattern of period two where highly transmissible and immune evasive strains repeat (Theorem 3.8). The most interesting case is when evolution exhibits chaotic behavior, making viral evolution unpredictable (Theorem 3.13). Our findings are illustrated by corresponding figures, charts and the bifurcation diagram (see Figs. 3.3a – 3.5).

## 2. Direction of the viral evolution: higher transmissibility or immune evasion?

In this section, after the introduction of the SIR model, we aim to ascertain the direction of the virus’s evolution. We seek to identify the circumstances under which the virus’s evolutionary path inclines toward heightened transmissibility or increased immune evasion. Additionally, we investigate the conditions under which a newly emerging strain can successfully invade the resident strain.

We consider an SIR model with recruitment rate of susceptibles *µ* > 0, and *µ* also equal the per capita death rate. Upon infection, susceptible individuals move to the infected compartment *I* at rate *βS*(*t*)*I*(*t*). Here, parameter *β* signifies the transmission rate of the resident strain. Subsequently, the infected individuals recover at rate *γ* > 0, leading them to transition to the recovered class *R*, who are immune to this strain. Therefore, we consider the following system of differential equations:

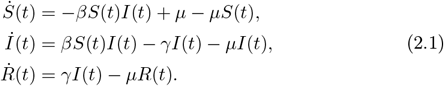

We consider a normalized population *N* = *S*(*t*) + *I*(*t*) + *R*(*t*) = 1, which is invariant for all *t*.

Now, by introducing invader strains (denoted by index *v*) into the system, we aim to discern whether the emergence of this invader strain is attributable to its capacity to evade immunity or its enhanced transmissibility. This introduction is made under the assumption that the system attains its endemic steady state:

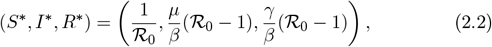

where 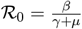 is the basic reproduction number. Note that the condition ℛ_0_ > 1 necessitates the relation *β* > *γ* + *µ*. This assumption ensures that the endemic steady state exists and is globally attractive.

In contrast to the slow mutation process inherent in many evolutionary models, our model distinguishes itself by allowing for the invasion of strains by a diverse array of variants, rather than solely mutants of the resident strain with very similar parameters. Many populations experienced such situation during COVID-19, where the evolution progressed to many different directions in other parts of the world, and countries were facing a large number of new strains originated from elsewhere, and some of those were able to invade and replace the previously dominant strain. These newly emerging strains, referred to as invader strains, differ from the resident strain in two distinct ways:

- Immune Evasion: Invader strains have the capability to evade immunity and infect individuals who have recovered from the resident strain (*R*). To quantify this, we introduce the parameter *p* ∈ [0, 1] which represents the fraction of recovered individuals from the resident strain that can be infected by the new strain.
- Transmissibility: Invader strains may exhibit either heightened or diminished contagiousness relative to the resident strain. This is delineated by the parameter *β*_*v*_ = *β* + *f* (*p*), where *f* (*p*) represents the trade-off function between transmissibility and immune evasion. A positive value of *f* (*p*) indicates an enhanced transmissibility of the invader strain compared to the resident strain, while a negative *f* (*p*) suggests a reduction in transmissibility. This trade-off function operates under the following assumptions:

(H1) *f* is continuously differentiable on [0, 1];

(H2) *f*′(*p*) *<* 0, for *p* ∈ [0, 1];

(H3) *f* (0) > 0 and *f* (1) *<* 0.

The early dynamics associated with the invader strain, represented by linearization at the endemic equilibrium, and characterized by the transmission rate *β*_*v*_, can be expressed as follows:

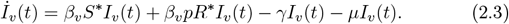

Therefore, the invasion reproduction number for the invader strains when the resident strain is in its endemic steady state, is

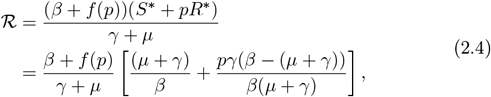

which denotes the number of secondary infections produced by an individual infected with the invasive variant over the course of their infectious period, within a population where resident strains have achieved equilibrium [20]. This can be interpreted as follows: An infectious individual remains infectious for 1*/*(*µ* + *γ*) time units. The number of new infections produced by a single infected host is given by (*β* + *f* (*p*))(*S*\* + *pR*\*) per unit time, since the available pool for infection with immune evasion parameter *p* is *S*\* + *pR*\*. The product of these two quantities is the number of secondary infected cases that are caused by one infection. The equilibrium abundance of uninfected hosts prior to the arrival of the infection compartment by invader strains is given by (2.2). Hence (2.4) represents the invasion reproductive ratio, which is a crucial quantity that determines whether a virus can spread in a host population. If ℛ *<* 1, the population infected by the invader is expected to diminish over time, whereas, ℛ > 1 measures the potential for the novel strain to continue spreading and potentially invade the population.

In the subsequent theorem, we will show that when *β* is small, the invasion reproduction number is a monotone decreasing function of *p*, indicating the emergence of a new strain with a higher transmission rate. Conversely, in case of high transmission rate *β*, circumventing the immune system is the most advantageous evolutionary strategy for the invader strain.

### Theorem 2.1

*Let f satisfy (H1)–(H3). We assume the resident strain of model* (2.1) *is in its endemic steady state, as given by* (2.2). *Then, there exists sufficiently small* δ > 0 *such that if β* ∈ (*µ* + *γ, µ* + *γ* + δ), *then the invasion reproduction number* ℛ (*p*) *decreases on* [0, 1], *and it attains its maximum at p* = 0 *and β*_*v*_ = *β* + *f* (0). *For large values of β*, ℛ *is an increasing function of p on* [0, 1], *hence the maximum of* ℛ (*p*) *occurs at p* = 1 *and β*_*v*_ = *β* + *f* (1).

*Proof*. To prove the theorem, it is enough to show that for small *β*, ℛ′(*p*) is negative, and for large *β*, ℛ′(*p*) is positive on [0, 1] (here′ denotes the derivative with respect to *p*).

Straightforward calculation yields

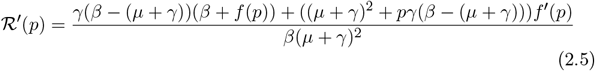

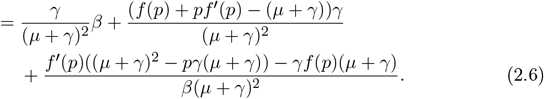

Since *f* ∈ *C*^1^[0, 1], there exist positive constants *k* and *K* such that |*f* (*p*)| *< K* and *K* ≤*f*′(*p*) ≤−*k* hold for all *p* ∈ [0, 1]. Combining this with (2.5), it follows that if *β* − (*γ* + *µ*) is sufficiently small, then ℛ′(*p*) *<* 0 holds for all *p* ∈ [0, 1], since with *β* →*µ* + *γ* we are left with *f*′(*p*)*/β <* −*k/β <* 0. This implies that the function ℛ is decreasing throughout and the maximum of ℛ occurs at *p* = 0.

On the other hand, considering the boundedness of both *f* and *f*′, we observe that for sufficiently large *β*, the first term dominates in (2.6), hence ℛ′(*p*) is positive for all *p* ∈ [0, 1] and it attains its maximum at 1.

The result of this theorem is intuitive: if the reproduction number is very large, then the recovered population at the steady state is also large, hence immune evasion can be very beneficial for a new strain. However, it does not gain much from immune evasion when the reproduction number is only slightly larger than one, since in this case the recovered population is small.

## 3. Analysis of the evolutionary process: global stability, periodicity, and chaos

Thus far, we have seen that a diminished transmission rate of the resident strain leads to the emergence of novel strains characterized by heightened transmissibility rather than increased immune evasion. Conversely, when *β* is large, the invading strains exhibit an increased capability to evade immunity.

In this section, we direct our attention towards identifying and closely examining the most invasive strain, i.e. the strain with the maximal invasion fitness. Therefore, we focus on answering the question of how this strain can maximize its reproduction number in presence of the resident strain.

To facilitate the mathematical analysis throughout the remainder of this paper, we employ a linear trade-off *f* with transmission advantage parameter *a*, and cost parameter *b* representing the expense of immunity evasion: *f* (*p*) = *a* − *bp*, where 0 *< a < b*.

In this case, the invasion reproductive number at the endemic steady state of the resident strain can be explicitly expressed as a function of *p*, and is characterized as a downward parabola of the form ℛ (*p*) = *Ap*^2^ + *Bp* + *C*, where

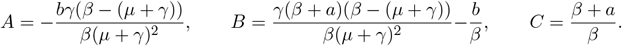

The maximum point of ℛ (*p*) is given by

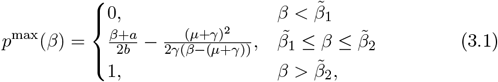

where

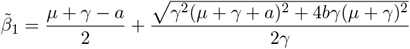

and

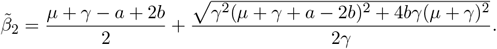

This new fittest strain is characterized by a novel transmission rate denoted as *β*_*v*_ = *β* + *a* − *bp*^max^, and from this point on, it takes the place of the resident strain in the system.

Fig. 3.1a presents a graphical representation of the invasion reproduction number ℛ (*p*) for three distinct values of the transmission rate *β*. Each value corresponds to distinct behaviors exhibited by the invasion reproduction function when its maximum value occurs at *p*^max^ = 0, *p*^max^ = 1, and *p*^max^ ∈ (0, 1), depicted as filled dots in the figure.

**Figure 3.1:**
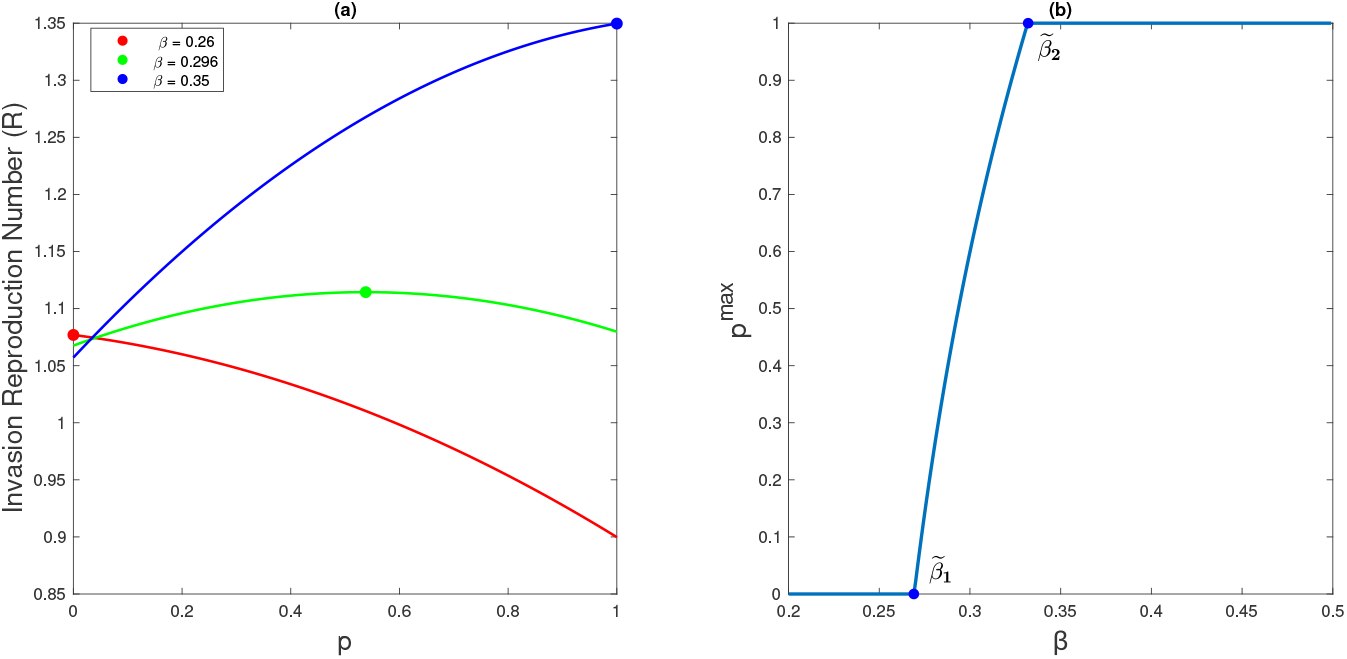
Panel (a) shows the invasion reproduction number with respect to *p* for three distinct values of the transmission rate parameter *β*, 0.26, 0.296, and 0.35 (curves red, green, and blue respectively). Each value corresponds to distinct behaviors exhibited by the invasion reproduction function when its maximum value occurs at *p*^max^ = 0, *p*^max^ = 1, and *p*^max^ ∈ (0, 1), respectively. Panel (b) shows the function *p*^max^(*β*) with respect to transmission rates of the resident strain. When the transmission rate *β* surpasses 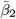, the maximum invasion reproduction number exhibits an increasing trend with maximum at *p* = 1 (*p*^max^ = 1), and for *β* close enough to *γ* +*µ, p*^max^ = 0. In both panels, *µ* = 3.5 · 10^−5^, *a* = 0.02, *b* = 0.1 and *γ* = 0.2.

By considering *p*^max^ as a function of *β*, the relationship between them is illustrated in Fig. 3.1b. This figure aligns with the implications of Theorem 2.1, indicating that as *β* increases, immune evasion becomes more favorable. Consequently, the invader strain is anticipated to propagate employing a strategy centered around immunity evasion, rather than exhibiting heightened transmissibility.

By iterating this procedure, we obtain a sequence of transmission rates driven by the difference equation

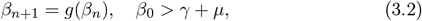

where *g* : (*γ* + *µ*, ∞) → (*γ* + *µ*, ∞),

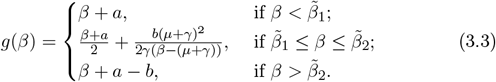

The following proposition contains some basic properties of the map *g*, which can be verified by straightforward calculation.

### Proposition 3.1.

*For any b* > *a* > 0, *function g defined in* (3.3),

i. *possesses a unique fixed point* 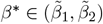 *defined as*

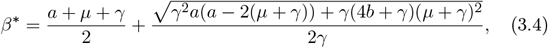
ii. *is linear and strictly increasing with constant slope* 1 *within the intervals* 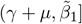 *and* 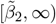,
iii. *is convex on* 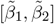, *moreover, it is strictly decreasing on* 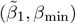 *and strictly increasing on* 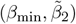, *where* 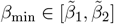 *is defined by*

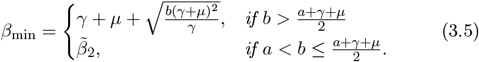

In the following subsections, we demonstrate a variety of behaviors, from stability, through periodicity, to chaos.

### 3.1 Global Convergence

Through the mathematical framework presented below, Theorem 3.2, we investigate the conditions under which transmissibility is stabilized, implying that over the long term, emerging variants will have approximately the same transmission rate. We illustrate this by providing explicit conditions under which the emergence of forthcoming strains, characterized by the sequence of transmission rates {*β*_*n*_}, converges to the fixed point.

#### Theorem 3.2.

*The unique fixed point β*\* *of the difference equation* (3.2) *is*

i. *locally asymptotically stable (LAS) if*

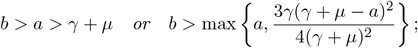
ii. *locally unstable if* 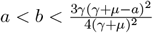 *and a* ≤ *γ* + *µ;*
iii. *globally asymptotically stable (GAS) (i*.*e. LAS and globally attractive) if*

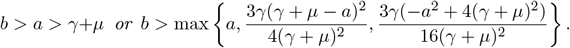

Before we prove Theorem 3.2, let us recall some results and tools related to local and global stability.

#### Theorem 3.3

([4, Theorem 2.1])

*If x*\* *is a fixed point of f* : (*c, d*) → (*c, d*), *then the fixed point is locally asymptotically stable if* |*f*′(*x*\*)| *<* 1, *and locally unstable if* |*f*′(*x*\*)| > 1.

#### Definition 3.4 ([12])

A function *ϕ*(*x*) envelops a function *f* (*x*) on the interval (*c, d*) if and only if for the unique fixed point *x*\*

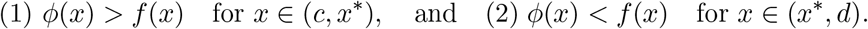

#### Theorem 3.5 ([13, Theorem 3])

*Assume that f maps the open interval* (*c, d*) *into itself. (The map f* (*x*) *may be discontinuous and/or multi-valued). Furthermore, assume that f* (*x*\*) = *x*\* *is the unique fixed point of f, and that there is a continuous self-inverse function ϕ*(*x*) *which envelops f* (*x*) *on* (*c, d*). *Then if x*_0_ *is any initial point and* {*x*_*n*_} *is any sequence consistent with x*_*t*+1_ = *f* (*x*_*t*_), *then* {*x*_*n*_} *converges to x*\*.

*Proof of Theorem 3*.*2*. (i) From Theorem 3.3, the local asymptotic stability of the function *g*(*β*) is established when the condition |*g*′(*β*\*)| *<* 1 is satisfied. An elementary calculation shows that if *a* > *γ* + *µ*, then *g*′(*β*\*) ∈ (0, 1). Moreover, if 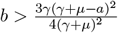 and *a* ≤ *γ* + *µ*, then *g*′(*β*\*) ∈ (−1, 0), concluding the proof of statement (i).

(ii)Similarly, when 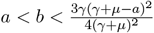 and *a* ≤ *γ* + *µ*, then *g*′(*β*\*) *<* −1. Thus, according to Theorem 3.3, the fixed point is locally unstable.

(iii)Assume that the condition of statement (iii) holds. To establish global stability, in view of Theorem 3.5 and statement (i), it suffices to construct a suitable function which envelopes the function *g*. To accomplish this, we introduce the self-inverse function *ϕ*(*β*) = 2*β*\* − *β*. We claim that *ϕ* is an enveloping function for *g* on (*γ* + *µ*, ∞). Without loss of generality, we show that *ϕ* envelopes function *g* over the interval 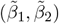 (since *ϕ* is strictly decreasing and 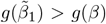 for 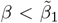, and 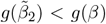, when 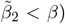. Given that the function *g* is convex on 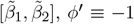, and *g*′(*β*\*) > −1, hence, to fulfill the first condition of Definition 3.4, it suffices to show that 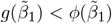. This inequality holds exactly when 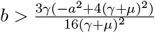.

Finally, this condition always holds when *b* > *a* > *γ* + *µ*, because 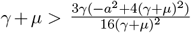. This completes the proof of statement (iii).

Fig. 3.2 shows that *g*(*β*) is enveloped by the linear function *ϕ*(*β*) = 2*β*\* − *β*. In this example, we have *a* > *µ* + *γ*, thus global asymptotic stability follows from Theorem 3.2.

Fig. 3.3a illustrates the bifurcation diagram of the function *g*(*β*) with *b* serving as the bifurcation parameter. This diagram reveals how varying the parameter *b* leads to changes in solutions of the difference equation (3.2), representing the sequence of transmission rates of emerging fittest strains. The pink and orange lines mark the critical points of *b* for local and global stability, respectively (under the condition *a < γ* +*µ*). Surpassing the threshold of local stability given by Theorem 3.2(i), solutions demonstrate convergence towards the fixed point. This means, over a sufficiently long period, all prevailing strains will practically have the same transmission rate. This is also illustrated in Fig. 3.4a, where any chosen initial point leads to a solution converging to this fixed point. These numerical evidences indicate that the notable gap between the mentioned thresholds (cf. statements (i) and (ii) of Theorem 3.2) could be filled. In other words, we conjecture that local asymptotic stability of the fixed point in fact implies its global stability. In the bifurcation diagram, a lower punishment (cost) parameter *b* is associated with increased complexity in behavior, as detailed in Sec. 3.3.

**Figure 3.2:**
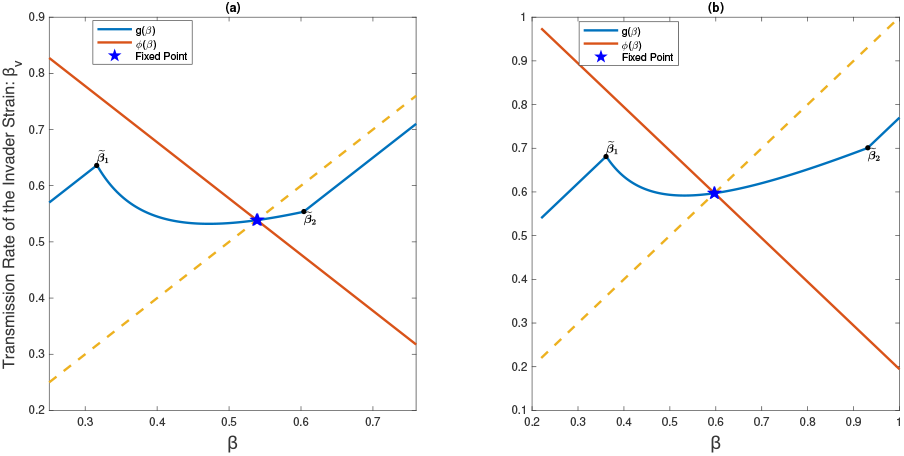
Graph of *g*(*β*) = *β* + *a*− *bp*^max^ plotted with the enveloping function *ϕ*(*β*) = 2*β*\* − *β* (red line) and identity line *β*_*v*_ = *β* (dashed line). In panel (a), the parameter *b* = *a*+0.05 is chosen such that 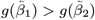, and in panel 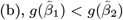 for *b* = 0.55. In this example, *a* = 1.6(*γ* + *µ*) > *µ* + *γ*, thus global asymptotic stability follows from Theorem 3.2. Other parameters are *γ* = 0.2, *µ* = 3.5 · 10^−5^.

**Figure 3.3:**
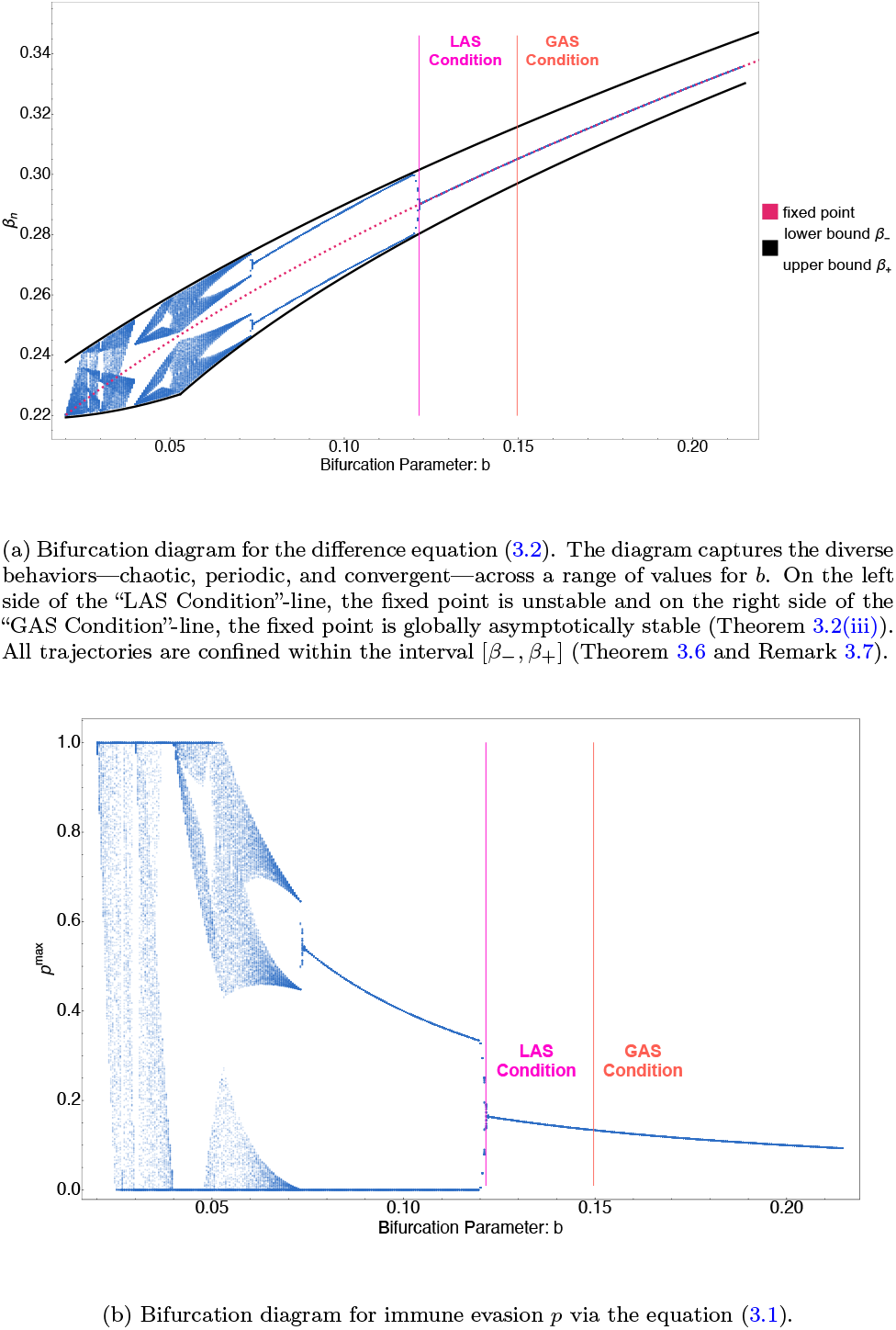
Bifurcation diagram for the equations (3.2) and (3.1). The system was iterated for *n* = 1000 steps from ten initial values for each *b* between 0.02 and 0.21 with step-size 0.0005, and the last 50 iterations are displayed in the plot. Here, *µ* = 3.5 · 10^−5^, *γ* = 0.2, and *a* = 0.02. (a) Cobweb plot for (3.2) with initial value β = 0.258 and parameter b = 0.155 tends to the unique fixed point. This diagram illustrates the case when the fixed point β* is globally asymptotically stable. (b) The cobweb plot with initial value β = 0.254 and b = 0.11 illustrates the manifestation of a two-cycle within the system. With multiple iterations, only two transmission rates are repeated alternately. (c) The cobweb plot, initialized with β = 0.2227 and parameter b = 0.03, reveals a 3-periodic solution, as also proved in Theorem 3.13.

**Figure 3.4:**
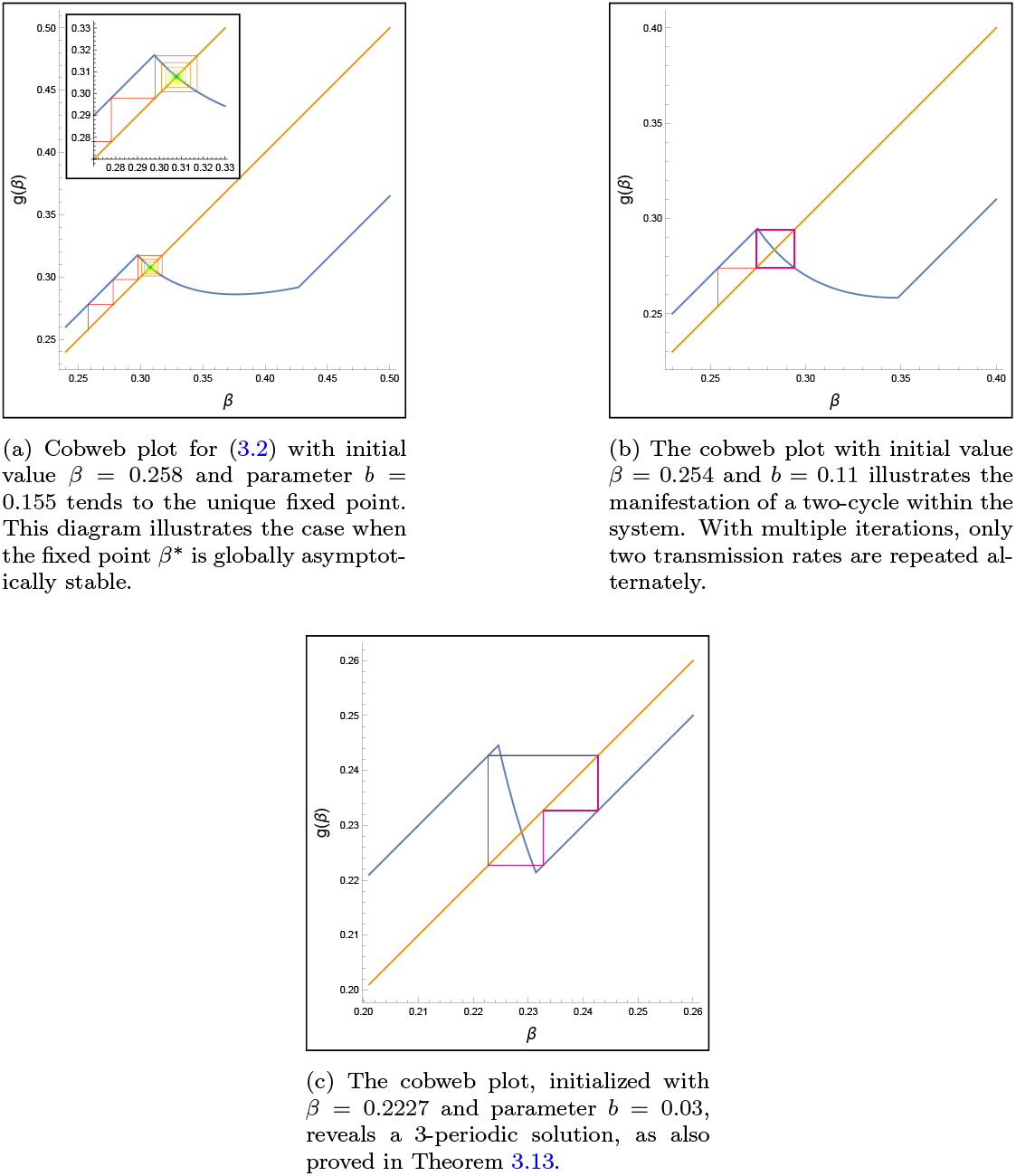
The cobweb plots correspond to three distinct behaviors of (3.2). *µ* = 3.5 10^−5^, *γ* = 0.2, and *a* = 0.02.

Figure 3.3b also illustrates the bifurcation diagram for the immune evasion parameter *p* via equation (3.1), with *b* serving as the bifurcation parameter. When *b* reaches a value where the system exhibits a two-periodic transmission rate, *p*^max^ also alternates between two values: *p*^max^ = 0 and some *p*^max^ ∈ (0, 1). This means that strains with a higher transmission rate will be followed by a strain having smaller transmission rate but with the ability to potentially reinfect a significant proportion of the recovered population (30–50% in this parameter region). Conversely, strains with lower *β* will be followed by *p*^max^ = 0, where the next strain will have the advantage of having a higher transmission rate, but in return, it will not be able to reinfect individuals recovered from the previous strain at all.

### 3.2 Attracting Interval

In the upcoming theorem, we identify an interval into which all solutions of the difference equation (3.2) enter and do not leave thereafter. This implies that, over an extended period, the transmission rates of the new strains remain within this specified interval.

#### Theore m 3.6.

*L et β*_+_ > sup *g* ((*γ* +*µ, β*\*] *and γ* +*µ < β*_−_ *<* inf *g* [*β*\*, *β*_+_]). *Then g*([*β*_−_, *β*_+_]) ⊆ [*β*_−_, *β*_+_], *and every solution of* (3.2) *enters* [*β*_−_, *β*_+_] *without leaving it again*.

*Proof*. This theorem is a direct application of Proposition 9.5 of [30].

#### Remark 3.7.

The supremum of the function *g* on (*γ* + *µ, β*\*] is easily observed to be 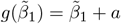. The determination of the infimum depends on the parameters and may manifest as 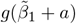, or *g*(*β*_min_), where *β*_min_ was defined in (3.5).

When 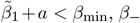 can be chosen arbitrarily from 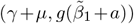. Conversely, for instances where 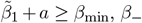 can be selected arbitrarily within the interval (*γ* + *µ, g*(*β*_min_)).

We depict the bounded interval, denoted as [*β*_−_, *β*_+_], within the bifurcation diagram shown in Fig. 3.3a. This illustration offers a visual confirmation that the transmission rates associated with the newly emerging strain will persist within this designated interval over an extended period.

### 3.3 Periodic Solutions and Complex Dynamics

In this section, we demonstrate that, in cases where the fixed point is locally unstable, there is at least one two-periodic solution. This is observed in the depicted solutions presented in Fig. 3.3a and 3.4b for specific values of the parameter *b*. These findings indicate that, despite multiple iterations and the emergence of numerous subsequent strains, only two transmission rates are repeated alternately in the system over the long term. In addition, within Theorem 3.13, we demonstrate that the difference equation (3.2), under specific conditions, exhibits chaotic behavior. This implies that the system’s dynamics is unpredictable, making it challenging to forecast whether the emergence of the next strain will be attributed to a heightened transmission rate or its capability to evade the immune system.

#### Theorem 3.8.

*If* 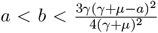 *and a* ≤ *γ* + *µ, then, there exists at least one two-periodic solution of the difference equation* (3.2) *different from β*\*.

To prove this theorem, we use the following concepts and results of [30].

#### Assumption 1

The function *f* : (*c*, ∞) → (*c*, ∞) is continuous, with a unique fixed point *x*\* > *c*, and is bounded on (*c*, ∞). Moreover, there exist *x*_1_ and *x*_2_, *c < x*_1_ *< x*\* *< x*_2_ *<* ∞, such that *f* (*x*_1_) > *x*_1_ and *f* (*x*_2_) *< x*_2_.

#### Theorem 3.9 ([30, Theorem 9.6])

*Assume that Assumption 1 holds and there is no fixed point of f* ^2^ *different from the unique fixed point x*\*. *Then all solutions of the difference equation x*_*n*+1_ = *f* (*x*_*n*_), *i*.*e*. {*x*_*n*_} *with n* ∈ N *with x*_0_ > *c, converge to x*\*, *as n* → ∞.

#### Definition 3.10.

For any *y* ∈ (*c*, ∞), let *M*_−_(*y*) represent the set of initial conditions *x* that are mapped onto *y* by the iterative application of the function *f*, i.e.

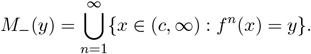

#### Proposition 3.11 ([30, Proposition 9.11])

*Assume that f* ^−1^({*x*}) *is countable for every x* ∈ (*c*, ∞). *Then M*_−_(*y*) *is countable for every point y* ∈ (*c*, ∞).

#### Remark 3.12.

Assumption 1, Theorem 3.9 and Definition 3.10 provided above, are found in [30] with *c* = 0. However, this can be easily addressed through a trivial change of variables.

Now, we are in position to prove Theorem 3.8.

*Proof of Theorem 3*.*8*. Considering Theorem 3.9 and the fulfillment of Assumption 1, it is adequate to demonstrate the existence of a solution that does not converge to *β*\*. According to Theorem 3.2, when 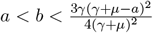 and *a* ≤*γ* +*µ, β*\* is locally unstable and *g*′(*β*\*) *<* −1. Since 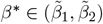, so *g*′ is continuous in a small neighborhood of the fixed point, therefore, there exists some *ϵ* ∈ (0, *β*\*−(*γ*+*µ*)] such that *g*′(*β*) *<* −1 for all *β* ∈ (*β*\*−*ϵ, β*\*+*ϵ*). Furthermore, since any point *β* ∈ (*γ*+*µ*, ∞) has a countable inverse set, more precisely, the cardinality of the set *g*^−1^({*β*}) is at most three (see Proposition 3.1 (ii)–(iii)), it follows that the set *M*_−_(*β*) is also countable as per Proposition 3.11. Consequently, there exists *β*_0_ ∈ (*γ* + *µ*, ∞), such that *β*_0_ ∉ *M*_−_(*β*\*). Assuming to the contrary that lim_*n*→∞_ *β*_*n*_ = *β*\*, there exists some *m* ∈ N such that |*β*_*n*_ − *β*\*| *< ϵ* for all *n* ≥ *m* and *β*_*n*_/= *β*\*, since *β*_0_ */ M*_−_(*β*\*). However, by the mean value theorem, for all *n* ≥ *m*, |*β*_*n*+1_ − *β*\*| = |*g*(*β*_*n*_) − *g*(*β*\*)| = |*g*(*ξ*_*n*_)| | *β*_*n*_ − *β*\*| >| *β*_*n*_ − *β*\*| > 0 holds for some *ξ*_*n*_ between *β*\* and *β*_*n*_. This contradicts the convergence of *β*_*n*_ towards *β*\*, proving our statements.

#### Theorem 3.13.

*Let* 0 *< a < b. If* 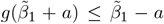 *or* 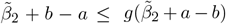 *holds, then the difference equation* (3.2) *is chaotic in the sense of Li and Yorke [17], i*.*e*.

1. *for every positive integer k there is a periodic point in* (*γ* +*µ*, ∞) *having period k;*
2. *there is an uncountable set S* ⊆ (*γ* + *µ*, ∞) *with no periodic points which satisfies the following conditions:*
  i. *for every p, q* ∈ *S with p*/= *q*,

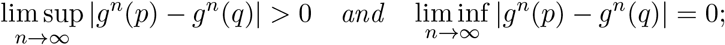
  ii. *for every p* ∈ *S and periodic point q* ∈ (*γ* + *µ*, ∞),

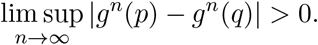

*Proof*. By virtue of [17, Theorem 1] it suffices to identify a *β*_0_ that fulfills condition *g*^3^(*β*_0_) ≤ *β*_0_ *< g*(*β*_0_) *< g*^2^(*β*_0_) or *g*^3^(*β*_0_) ≥ *β*_0_ > *g*(*β*_0_) > *g*^2^(*β*_0_).

When 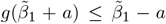, then let 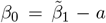. Note that, since *g* maps the interval (*γ* + *µ*, ∞) into itself, *γ* + *µ < β*_0_ must hold. Then the inequalities

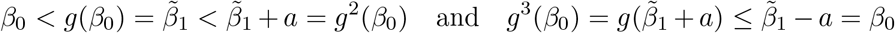

clearly hold, so *g*^3^(*β*_0_) ≤ *β*_0_ *< g*(*β*_0_) *< g*^2^(*β*_0_) is satisfied.

Similarly, if 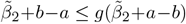 is fulfilled, then setting 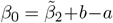 leads to

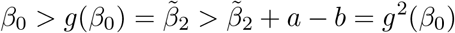

and

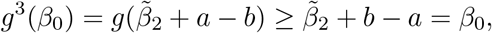

that is, *g*^3^(*β*_0_) ≥ *β*_0_ > *g*(*β*_0_) > *g*^2^(*β*_0_) holds, just as required.

#### Corollary 3.14.

*If either* 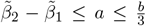 *or* 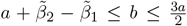 *is satisfied, then the difference equation* (3.2) *is chaotic in the sense of Li and Yorke*.

*Proof*. Assuming 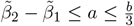 yields 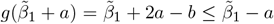 while condition 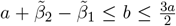 immediately implies 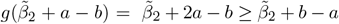. Hence, conditions of Theorem 3.13 are satisfied in both cases.

A straightforward implication of Theorem 3.13 is the existence of a three-periodic solution. Fig. 3.4c offers numerical validation of this phenomenon, presenting evidence that corroborates the theoretical predictions made by the theorem.

Fig. 3.5 illustrates four distinct regions, each representing a different dynamical behavior of the equation (3.2). The blue and yellow regions denote the domains where the fixed point is globally and locally asymptotically stable, respectively, whereas the green region highlights areas of instability where 2-periodic orbits emerge, as delineated in Theorems 3.2 and 3.8. The gray region (obtained in Theorem 3.13 (gray) and Corollary 3.14 (light gray region)) reflects the chaotic aspects of the difference equation (3.2), within which predicting the behavior of subsequent strains becomes challenging.

**Figure 3.5:**
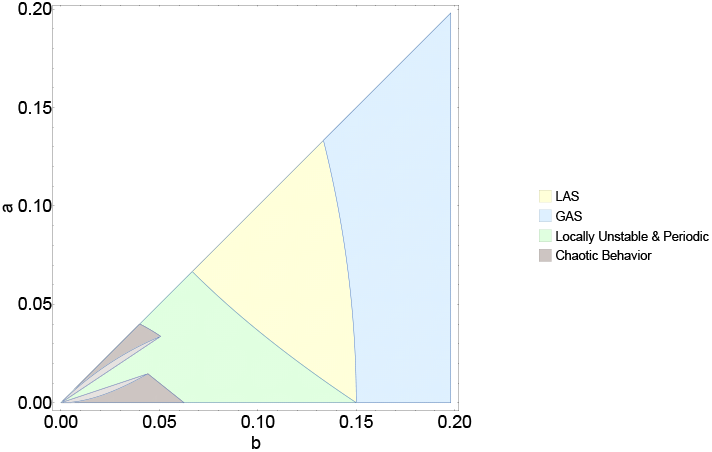
The figure delineates four distinct regions, each corresponding to a unique dynamical behavior of the system as defined by the difference equation (3.2). The regions colored in blue and yellow represent domains where the fixed point exhibits global and local asymptotic stability, respectively. The green regions are zones of instability, as given in Theorem 3.2. The area depicted in gray encapsulates chaotic dynamics of (3.2). Here *γ* = 0.2 and *µ* = 3.5 · 10^−5^. (a) Stable transmission rates yield similar waves and the same prevalence at steady state (shown in the inset) across subsequent strains. Here *b* = 0.208, *β* = 0.404, and *a* = 0.2

Figure 3.6 illustrates the prevalence of infection for each strain within the context of the presented model (2.1), following the scheme presented in Figure 1.1. This illustrative figure was constructed by integrating the SIR model, and when the solution approached the endemic steady state, we switched to a new strain (i.e., new *β*) selected by maximizing invasion fitness via the trade-off. Depending on the immune evasion parameter *p* of the new strain, we reallocated recovered and susceptible individuals to create the initial state for the next period. Then we repeated the procedure by newer and newer strains.

**Figure 3.6:**
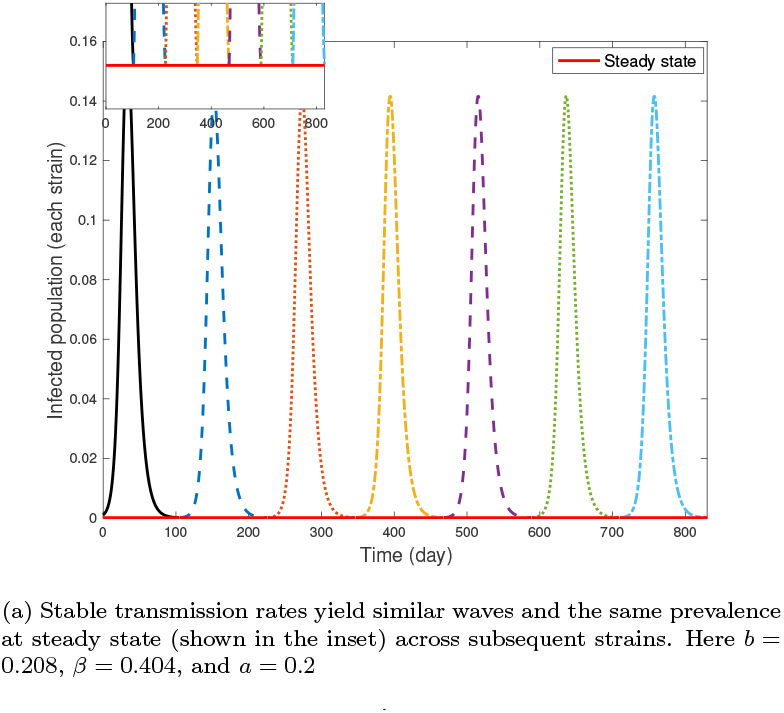
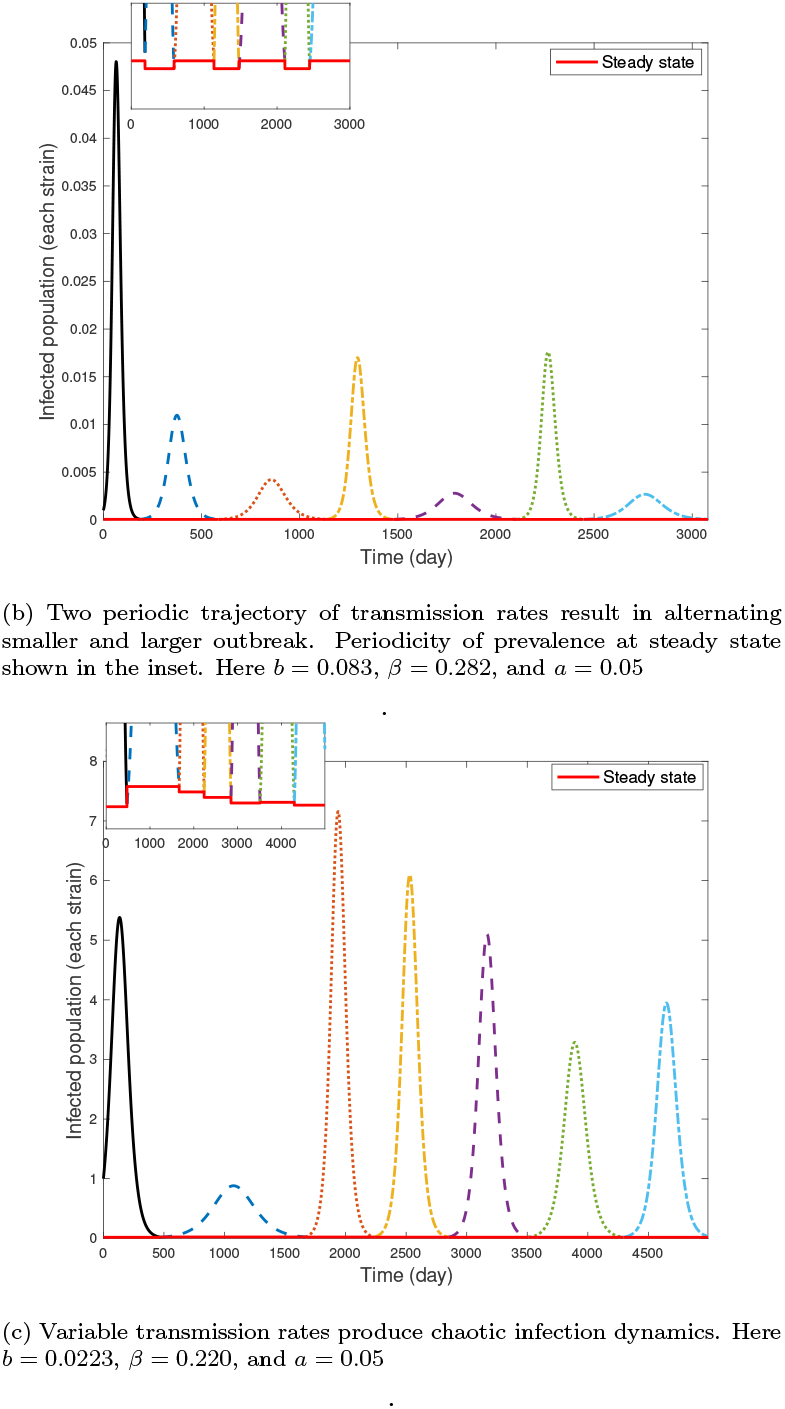
Prevalence dynamics of infection under varying transmission rate scenarios. The initial value of *S, I*, and *R* are 0.999, 0.001, and 0 respectively; *µ* = 3.5· 10^−5^, *γ* = 0.2.

Panel (a) corresponds to the conditions outlined in Theorem 3.2, where the transmission rates of strains converge to an equilibrium. Under these conditions, a stable transmission rate ensures a uniform steady-state prevalence across all strains. As observed, the infection levels for all strains remain consistent over time, showing a stable and predictable pattern of infection dynamics.

Panel (b) corresponds to a stable 2-periodic scenario. After the first wave, we experience a fast convergence in the trajectory of the dynamics of transmission rates to a 2-periodic pattern. This periodicity in transmissibility leads to alternating prevalence levels. Higher prevalence corresponds to a greater proportion of the infected population, indicating a more harmful evolutionary outcome.

In panel (c) each strain possesses a different transmission rate, resulting in a highly irregular and unpredictable pattern of infection prevalence. This behavior corresponds to the chaotic regime of the difference equation (3.2). The figure highlights how variations in transmissibility across strains can drive complex outcomes in infection prevalence, making it challenging to predict long-term dynamics.

## 4. Discussion

Gaining insights into the evolution of pathogens’ traits is significant not only in the context of evolutionary dynamics but also holds crucial implications for epidemiology and public health. These evolutionary traits include virulence, transmission efficiency, replication rate, resistance to stress and antimicrobials, etc. Mechanisms that allow pathogens to avoid, suppress, or manipulate the host’s immune system are also vital for their long term survival. Understanding the evolution of host immune evasion is central to designing influenza vaccines, since influenza strains in subsequent years may evade prior immunity [9, 14], and this issue gained attention during the COVID-19 pandemic as well [28].

The evolution of these traits is often intertwined via various trade-offs. Most notably, the trade-off between virulence and transmissibility has been investigated in numerous research papers. In comparison, the immune evasion-transmissibility trade-off received less attention, and has not been explicitly considered in mathematical models of viral evolution. By incorporating this trade-off into a novel evolutionary model, we explored the emergence of new variants and their long term evolutionary patterns.

Our starting point was an epidemiological model (2.1) considering a single resident strain. Novel strains emerge with a transmission rate denoted by *β*_*v*_ = *β* +*f* (*p*), where *p* ∈ [0, 1] is the immune-evasion, and *f* (·) is a decreasing function expressing the trade-off. Strains with *p* = 0 lack the ability to evade the immune response in individuals who have acquired immunity and currently belong to the recovered (R) compartment. Conversely, strains with *p* = 1 are capable of infecting all individuals who have recovered from the previous strain. As demonstrated in Theorem 2.1, when the transmission rate of the resident strain, *β*, is sufficiently close to *γ* + *µ*, the invasion reproduction number ℛ (*p*) decreases across the range of *p*. Consequently, the new strain exhibits a higher transmission rate, and invasion fitness is maximized at *p* = 0. Conversely, for cases where *β* ≫ (*γ* + *µ*), immune evasion becomes advantageous and invasion fitness is maximized at *p* = 1. For intermediate values of *β*, invasion fitness is maximized at some 0 *< p <* 1.

We assume that evolution selects the most invasive strain (with maximal invasion reproduction number), characterized by a transmission rate *β*_*v*_ = *β* + *f* (*p*^max^). This new variant replaces the resident strain, and we let the system into the new endemic steady state, initiating a cycle of strain replacements. This process can be reduced mathematically to a difference equation (see (3.2)). For the sake of analytic results, we focused on a linear trade-off function *f* (*p*) = *a*−*bp*. Our findings reveal that this methodological simplification does not compromise the complexity of the observed dynamics, which range from the global stability of the fixed point to the emergence of periodic and even chaotic behavior.

To find conditions for the global stability of the unique fixed point of (3.2), we applied the enveloping technique of [12]. Our investigation uncovered that, within a specific parameter range for *b*, the fixed point is globally asymptotic stable. This suggests that over time, new variants consistently exhibit similar intrinsic transmission rates of infection. Additionally, our analysis has demonstrated that the emergence of instability is accompanied by both periodic and chaotic behaviors. Specifically, Theorem 3.6 establishes the existence of at least one two-periodic solution, suggesting that the system can alternate between only two transmission rates over extended periods. Theorem 3.13 delineates the conditions under which the solutions of the difference equation (3.2) exhibit chaotic behavior. This chaotic nature renders the system’s dynamics inherently unpredictable in the long run. These diverse behaviors are visually represented in Fig. 3.5, where each colored region corresponds to one of the following dynamics of the equation (3.2): GAS, LAS, locally unstable, and chaotic/periodic. Bifurcation diagram 3.3a and cobweb plot 3.4 offer a visualization of the evolution of the given model across successive iterations, depicting its dynamics from an initial value. In Fig. 3.4a (and a segment of Fig. 3.3a), the convergence to the fixed point is evident, however it is not universal. When the conditions outlined in Theorem 3.2 are not met, the system may exhibit periodic and even chaotic behaviors, as depicted in Figs. 3.4b, 3.4c, and the left side of Fig. 3.3a, for 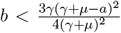. In the bifurcation diagram, one can see that reducing the cost parameter *b* is directly correlated with an escalation in behavioral complexity. This dynamics fosters a more unpredictable and complex pattern of viral evolution. As it was observed during the pandemic, SARS-CoV-2 viral evolution is inherently unpredictable [11]. Our results offer a mechanism via a simple trade-off that naturally leads to such unpredictability.

Naturally, there are several limitations to this approach. To focus on the impact of the transmissibility/immune evasion trade-off, we ignored other complex phenomena that influence the disease transmission dynamics, and restricted ourselves to a simple SIR model. We assumed that the most invasive strain replaces the resident strain. This is a strong assumption which simplifies our framework, but it might not capture the full range of potential evolutionary behaviors. For instance, relaxing this assumption could lead to scenarios where multiple strains co-circulate with significant prevalences, or where the success of invasion is influenced by factors such as host immune landscapes, heterogeneity in immunity in the population, or other selection pressures. Additionally, we ignored superinfection, competition between multiple strains, or immune memory from previous infections, which could alter the observed dynamics. However, in the context of SARS-CoV-2 evolution during the COVID-19 pandemic, in the first few waves we have not observed significant coexistence of strains. Instead, new variants have quickly and consistently outcompeted the previous strains and established dominance, as seen with the successive emergence of variants like Alpha and Delta. This offers that, at least for this particular scenario, the evolutionary trajectory aligns with the dominance of the fittest strain, possibly driven by its higher transmissibility or immune evasion capabilities. Nevertheless, our results point out that assuming such simple mechanisms and even a linear trade-off can lead to complicated dynamics.

Predicting the emergence of new viral strains is a major challenge. Our results indicate that even under a simplified model, where we assume a linear trade-off between transmissibility and immune evasion, the dynamics of strain emergence can be highly unpredictable and even chaotic in certain conditions. This suggests that, despite the simplicity of our model, the emergence of new strains can exhibit complex and erratic behaviors for a range of pathogens as well. Such findings highlight the inherent unpredictability in viral evolution, which is especially relevant when considering real-world scenarios, such as the evolution of SARS-CoV-2 during the pandemic.

Future research may consider more complicated disease dynamics models, in particular, our method is applicable to any compartmental model (which may include even differential delay equations) that has a unique, globally asymptotically stable endemic equilibrium. It would also be interesting to study other functional forms of trade-off, or multiple trade-offs between various traits. This way one can potentially assess the robustness of these results as well as gain further insight into viral evolution. Furthermore, the bifurcation diagram in Fig. 3.3a suggests that our result on global stability in Theorem 3.2 is not sharp. A potential avenue for future research is the application of different enveloping functions, which may result in sharper conditions for global stability. We conjecture that local stability of the equilibrium of the difference equation always implies its global stability as well.

## Acknowledgement

We are grateful to the anonymous reviewers for their insightful feedback and suggestions which have greatly improved the quality and clarity of this paper.

G. S. was supported by Marie Skłodowska-Curie EvoGamesPlus grant no. 955708. Á. G. was supported by NRDI Funds TKP2021-NVA-09, ÚNKP-23-5, NKFIH-FK 142891 and by the János Bolyai Research Scholarship of the Hungarian Academy of Sciences.. G. R. was supported by KKP 129877, RRF-2.3.1-21-2022-00006, TKP-2021-EGA-05.

